# Sp100A isoform promotes HIRA histone chaperone localization to PML nuclear bodies

**DOI:** 10.1101/2025.03.06.641437

**Authors:** Ashley N. Della Fera, Warda Arman, Maceo E. Powers, Alix Warburton, Alison A. McBride

**Affiliations:** Laboratory of Viral Diseases, National Institute of Allergy and Infectious Diseases, National Institutes of Health, Bethesda, Maryland, USA

**Keywords:** PML-NBs, Sp100, HIRA, histone chaperone, SUMO, keratinocyte

## Abstract

PML nuclear bodies (PML-NBs) are dynamic subnuclear structures important for chromatin dynamics and anti-viral defense. In this study we investigate the role of Sp100 isoforms in promoting localization of the H3.3 histone chaperone HIRA to PML-NBs in human keratinocytes. Sp100 knockout (KO) cell lines were generated using CRISPR-Cas9 technology and shown to display normal keratinocyte differentiation and PML-NB formation. However, HIRA and its associated complex members (UBN1 and ASF1a) failed to localize to PML-NBs in the absence of Sp100, even after interferon stimulation. Exogenous expression of the four main isoforms of Sp100 showed that the Sp100A isoform is the primary driver of HIRA localization to PML-NBs, with the SUMO interacting motif (SIM) playing an important role. These findings highlight the functional diversity of the Sp100 isoforms in modulating chromatin dynamics at PML-NBs.

## Introduction

Promyelocytic leukemia (PML) nuclear bodies (PML-NBs) recruit various cellular proteins and are comprised of numerous constitutively and transiently associated proteins (Van Damme *et al*, 2010) that function in a vast array of cellular processes, including, but not limited to, senescence, regulation of chromatin structure and gene expression, the ubiquitin pathway, the DNA damage response, and the innate immune response to viral infection (Scherer & Stamminger, 2016). Various chromatin-related modulators, such as histone modifiers, histone readers, and histone chaperones localize to PML-NBs, which reflects their role in chromatin structure and dynamics (Corpet *et al*, 2020). The replication-independent histone variant H3.3 is also enriched in PML-NBs and this is facilitated by two histone chaperone complexes, Daxx/ATRX and HIRA/UBN-1/CABIN1 (Choi *et al*, 2024). Daxx is a constitutive PML-NB protein that complexes with the chromatin remodeler a-thalassemia/mental retardation syndrome protein (ATRX) to aid in the maintenance of heterochromatin (Goldberg *et al*, 2010) and also to maintain a soluble non-nucleosomal pool of histone H3.3 at PML-NBs (Delbarre *et al*, 2013; Drane *et al*, 2010). HIRA primarily mediates H3.3 deposition in regions of the chromatin that are transcriptionally active (Goldberg *et al*., 2010; Placek *et al*, 2009; Zhang *et al*, 2017), but HIRA has also been implicated in chromatin silencing (Sherwood *et al*, 1993; van der Heijden *et al*, 2007). Daxx is constitutively associated with PML-NBs, while the histone H3.3 chaperone HIRA only associates with PML-NBs in response to cellular senescence (Banumathy *et al*, 2009; Ye *et al*, 2007; Zhang *et al*, 2005), interferon (IFN) stimulation, and viral infection (Cohen *et al*, 2018; Kleijwegt *et al*, 2023; McFarlane *et al*, 2019; Rai *et al*, 2017).

IFN induced localization of HIRA to PML-NBs depends on the Sp100 protein, which is one of the major components of PML-NBs (Kleijwegt *et al*., 2023; McFarlane *et al*., 2019). As shown in Figure 1A, Sp100 has four main isoforms, Sp100 A, B, C and HMG (Fraschilla & Jeffrey, 2020; Negorev *et al*, 2009) that share an N-terminal region, which contains a homogeneously staining region (HSR) domain, a SUMOylation site and a SUMO interaction motif (SIM) (Seeler *et al*, 1998). The three longer isoforms also contain a SAND (Sp100, Aire-1, NucP41/75, DEAF-1) DNA-binding domain (Bottomley *et al*, 2001). Sp100C contains an additional bromodomain and plant homeodomain (PHD) (Zhang *et al*, 2016), while Sp100HMG contains two high mobility group boxes (Seeler *et al*., 1998). Evaluations of the effects of the various Sp100 isoforms on viral and cellular gene expression have identified both activating and repressive functions; Sp100A primarily potentiates transcription (Berscheminski *et al*, 2014; Newhart *et al*, 2013; Wasylyk *et al*, 2002), whereas Sp100B, Sp100C, and Sp100HMG are repressive (Isaac *et al*, 2006; Ishov *et al*, 1999; Negorev *et al*., 2009; Wilcox *et al*, 2005). For example, Sp100A is retained in active PML-associated adenoviral replication centers, while the SAND domain containing isoforms were displaced (Berscheminski *et al*., 2014). Our laboratory has also shown that the three longer SAND domain containing isoforms repress HPV transcription and replication (Stepp *et al*, 2013). We hypothesize that the transcriptionally activating and repressive functions of the various Sp100 isoforms could influence the localization of HIRA to PML-NBs.

**Figure 1.**
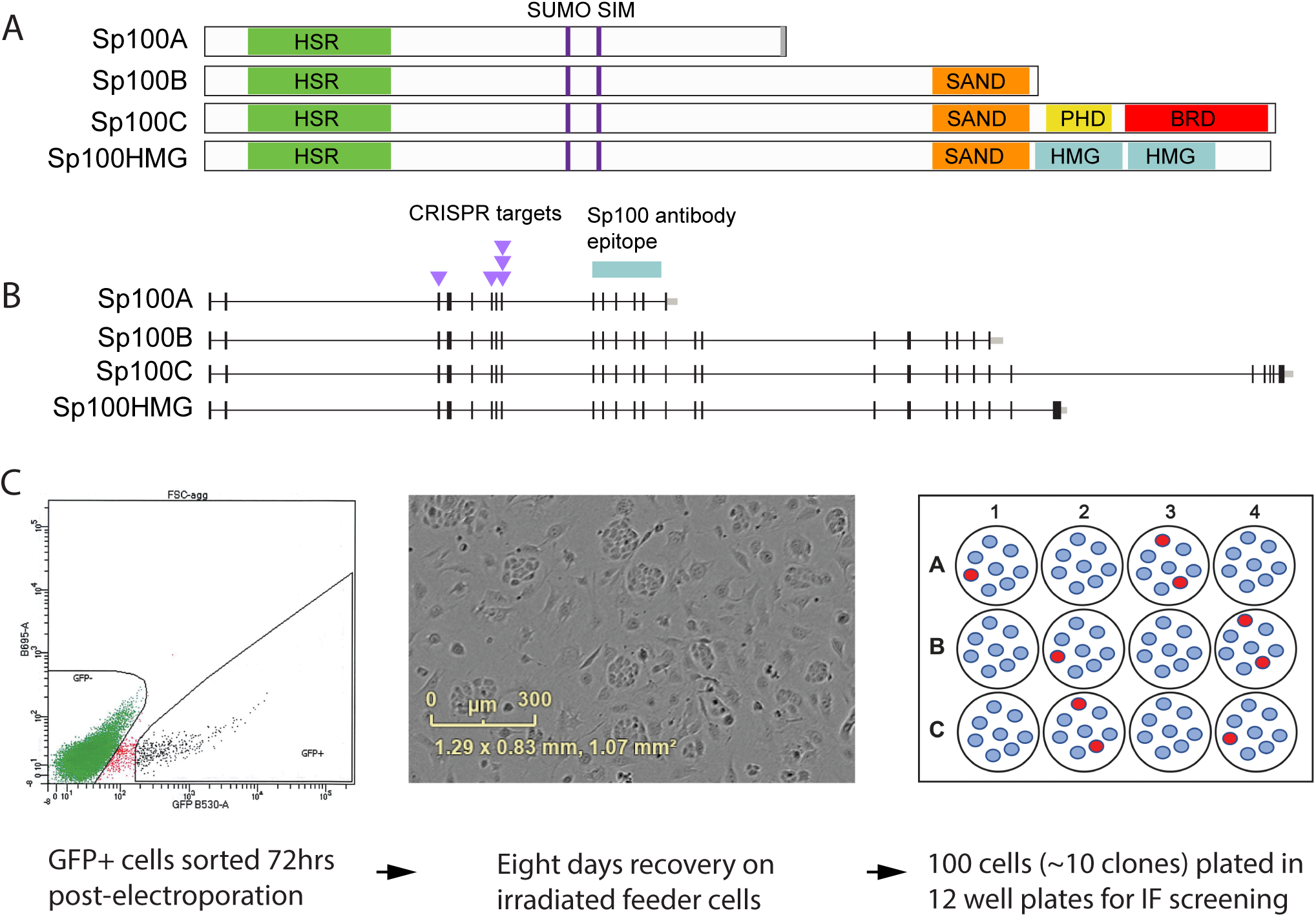
Generation of Sp100 Knockout NIKS by CRISPR. **A.** Map of the four major isoforms of Sp100. Indicated are the homogeneously staining region (HSR) domain, a SUMOylation site and a SUMO interaction motif (SIM), the SAND (Sp100, Aire-1, NucP41/75, DEAF-1) DNA-binding domain, the bromodomain and plant homeodomain (PHD) and high mobility group domains (HMG). **B.** Exon map of the four major isoforms of Sp100. The CRISPR guide RNAs were designed to target exons 3, 6 and 8. The polypeptide immunogen shown was used to generate the Sp100 antisera used for screening. **C.** Strategy to isolate NIKS Sp100 KO cells. Five plasmids encoding a GFP protein and each containing a guide RNA targeting So100 were electroporated into NIKS cells. Seventy-two hours later, GFP positive cells were isolated by fluorescent activated cell sorting and plated onto irradiated J2 3T3 feeders to recover. After eight days, 100 cells were plated into each well of multiple 12-well dishes.

In this study, we analyzed the interplay between the histone H3.3 chromatin assembly pathway and PML-NBs. Specifically, we investigated the role of Sp100 in promoting the association of HIRA with PML-NBs. We show that HIRA localization to PML-NBs is dependent on the Sp100A isoform and that the SIM motif of Sp100A promotes this association.

## Results

### Generation of Sp100 deficient NIKS cell lines

NIKS is a spontaneously immortalized keratinocyte cell line that has retained the ability to terminally differentiate into a stratified epithelium and is often used in the study of HPV infection (Allen-Hoffmann *et al*, 2000) and has been used to generate human skin grafts. To study the role of Sp100 in keratinocyte biology, we generated Sp100 knockout (KO) NIKS cell lines using CRISPR technology. We designed five CRISPR Cas9 guide RNAs to generate deletions in exons 3, 6 and 8 of the Sp100 gene (Figure 1B). The guide sequences were cloned into the PX458 plasmid, which also expresses GFP, and the resulting plasmids were electroporated into NIKs cells. As outlined in Figure 1C, seventy-two hours after transfection, GFP positive cells were selected by flow cytometry and plated onto irradiated feeder plates to allow recovery and expansion of the cells. NIKS cells have a plating efficiency of ∼10%, and so we plated 100 cells into each well in multiple 12-well dishes with the expectation of ∼ ten clones per well, which would allow effective screening for knockout clones by immunofluorescence. The Sp100 antibody used to confirm that Sp100 protein expression was eliminated recognizes all four isoforms and was raised against amino acids 312-411 of Sp100A (encoded by exons 9 to 13). After an initial screen of 36 clones by combined PML and Sp100 immunofluorescence, seven clones were expanded for characterization. These consisted of five Sp100 knockout clones with no Sp100 expression (Sp100 KO 1-5), and two clones derived from control cells (Vector 1-2) that had been transfected with the parental PX458 plasmid and underwent the same sorting and selection procedures. These clones were expanded into cell lines and the absence of Sp100 was verified by immunofluorescence (Supplementary Figure 1A) and Western blot (Supplementary Figure 1B).

### Genotype of the NIKS Sp100 KO clones

To genotype the cell clones, the regions targeted by CRISPR were PCR amplified, ligated into the PCR2.1 TOPO cloning vector and analyzed by Sanger sequencing. A minimum of eight clones were selected per exon for each cell line. As shown in Supplementary Table 1, each KO clone had mutations in exon 3 and exon 8, but not in exon 6. Sp100 KO 1, 2 and 3 cell lines harbored various deletions around the guide RNA that targeted exon 3, and also contained 35bp and 36bp deletions in exon 8. Sp100 KO 4 and 5 had an identical 158bp insertion at the guide site in exon 3 that is derived from 28S rRNA. These cells also had additional 35bp and 36bp deletions in exon 8, similar those in KO cells 1, 2 and 3.

### The absence of Sp100 does not affect keratinocyte differentiation

We have previously analyzed the role of Sp100 in HPV31 containing CIN612 9E keratinocytes using siRNA technology and observed an approximate three-fold increase in the late differentiation marker filaggrin after calcium induced differentiation (Stepp *et al*, 2017). To determine if the NIKS Sp100 KO cells can differentiate normally into a stratified epithelium, we generated organotypic rafts from the NIKS vector and Sp100 KO cells. As shown in Supplementary Figure 2, the SP100 KO cells formed a stratified epithelium that was indistinguishable from that formed by vector control or parental NIKS cells. We conclude that stable knockout of this protein does not affect keratinocyte differentiation.

### PML-NB formation and localization of Daxx are unaltered in Sp100 KO keratinocytes

Our laboratory has shown previously that transient downregulation of Sp100 siRNA does not alter PML-NB formation, or localization of the PML-NB resident protein Daxx, in primary HFKs (Stepp *et al*., 2013). To confirm that this is also the case in the NIKS Sp100 KO cells, NIKS parental, vector control, and Sp100 KO cells were treated with or without type I IFN and stained with antibodies against PML, Sp100, and Daxx and visualized by confocal microscopy. Representative images of parental NIKS, vector 2 and Sp100 KO3 cells are shown in Figure 2A and quantitation of all cell lines in Figure 2B and 2C. PML and Sp100 were highly responsive to stimulation with IFNα in NIKS parental and control cells; PML and Sp100 containing nuclear bodies were more intense and increased in number. Sp100 and Daxx were almost always associated with PML-NBs in NIKS parental and control cells regardless of stimulation with IFN (Figure 2A). However, while PML-NBs were completely devoid of Sp100 in the KO cells, both the average number of PML-NBs per nuclei (Figure 2B) and percentage of PML-NBs per nuclei containing Daxx (Figure 2C) were similar to those detected in the parental and control cells. Taken together, these data demonstrate that PML-NB formation and association with Daxx do not require Sp100 proteins.

**Figure 2.**
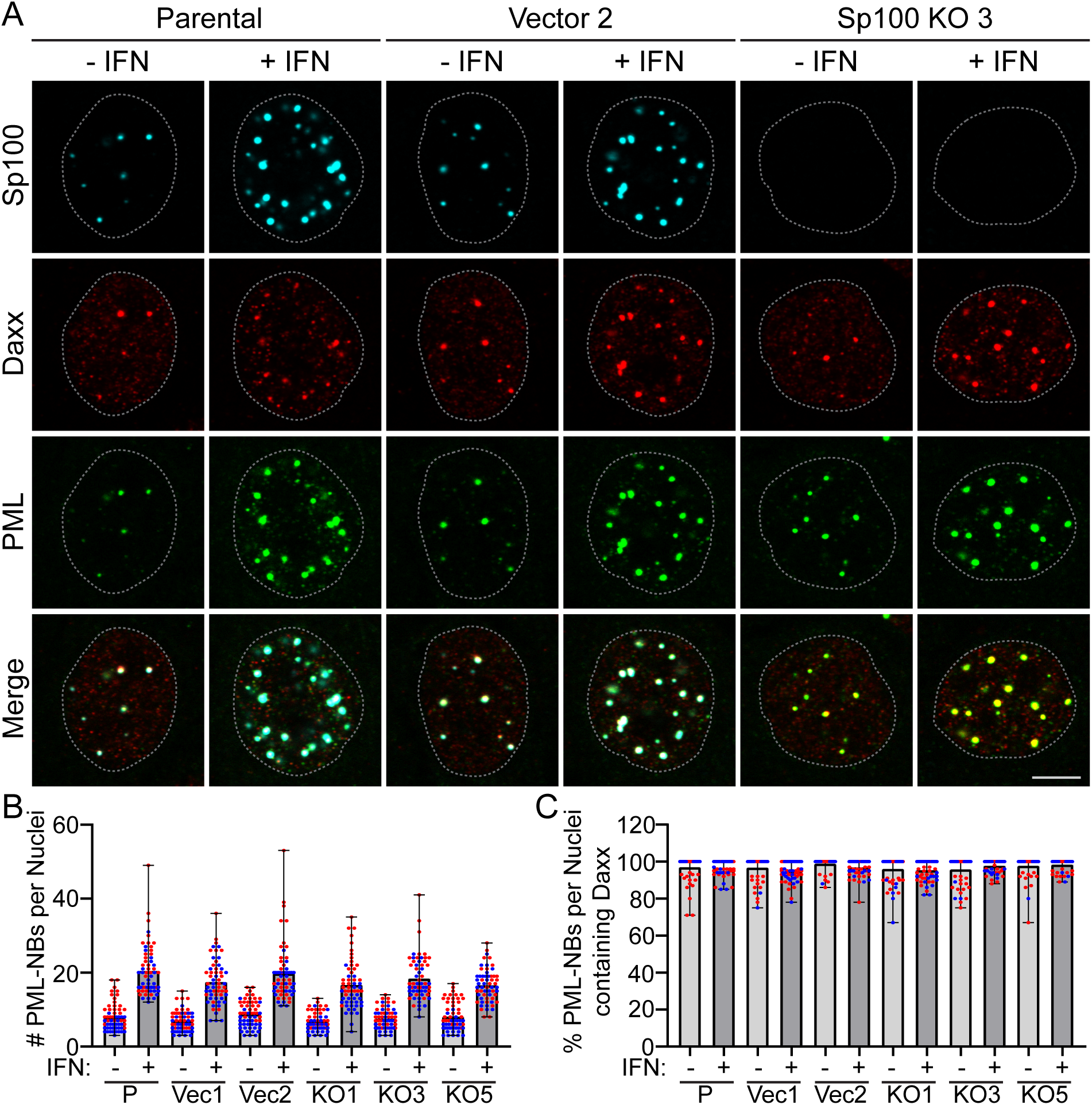
PML-NB formation and Daxx localization are unaltered in Sp100 KO keratinocytes. **A.** Representative immunofluorescence staining of PML (pseudo-colored green), Daxx (pseudo-colored red), and Sp100 (cyan) in NIKS Parental, Control Vector 2, and Sp100 KO3 cells with or without IFNα treatment for 48 hours. The white dotted lines outline the nuclei as defined by DAPI staining. Scale bar, 5 μm. **B.** The number of PML-NBs per nuclei. **C.** The percentage of PML-NBs per nuclei containing Daxx. For panel B and C quantitation, in NIKS Parental [P], Vector [Vec] 1 and 2, and Sp100 KO clones 1, 3, and 5; manually scored 30 nuclei in each of two independent experiments. The mean is shown, along with the individual nuclei (red and blue circles denote nuclei from two independent experiments); Error bars represent the range of all values.

To assess whether the levels of the PML or Daxx proteins changed in the NIKS Sp100 KO cells, protein lysates prepared from NIKS, vector, and Sp100 KO cells with or without stimulation with IFNα were analyzed by immunoblot (Supplementary Figure 3). As expected, PML and Sp100 protein levels were increased following stimulation with IFN in NIKS parental and control cells and Sp100 proteins were not detected in the NIKS Sp100 KO cells in either condition. In the Sp100 KO clones, the protein levels of PML and DAXX were similar to those observed in NIKS parental and vector cells. Thus, the protein levels of PML and Daxx are unaltered in the absence of Sp100.

### Sp100 is required for localization of HIRA to PML-NBs

The histone chaperone HIRA associates with PML-NBs in response to IFN, in a cell line specific manner, and McFarlane et al. and Kleijwegt et al. report that this is dependent on Sp100 (Kleijwegt *et al*., 2023; McFarlane *et al*., 2019). To determine whether IFN induced localization of HIRA to PML-NBs also occurs in NIKS keratinocytes, and whether Sp100 is required for this localization, we assessed the colocalization of HIRA with PML-NBs in NIKS, control, and Sp100 KO cells before and after stimulation with IFN. All NIKS control and Sp100 KO clones were assessed and are represented in Figure 3 by NIKS control 2 and Sp100 KO3 cells. As shown in Figure 3A, minimal amounts of HIRA were detected in a few PML-NBs in ∼20% cells, however this association increased greatly after IFN treatment (Figure 3B). In contrast, HIRA was seldom associated with PML-NBs in NIKS Sp100 KO cells either in the absence or presence of IFN (Figure 3B).

**Figure 3.**
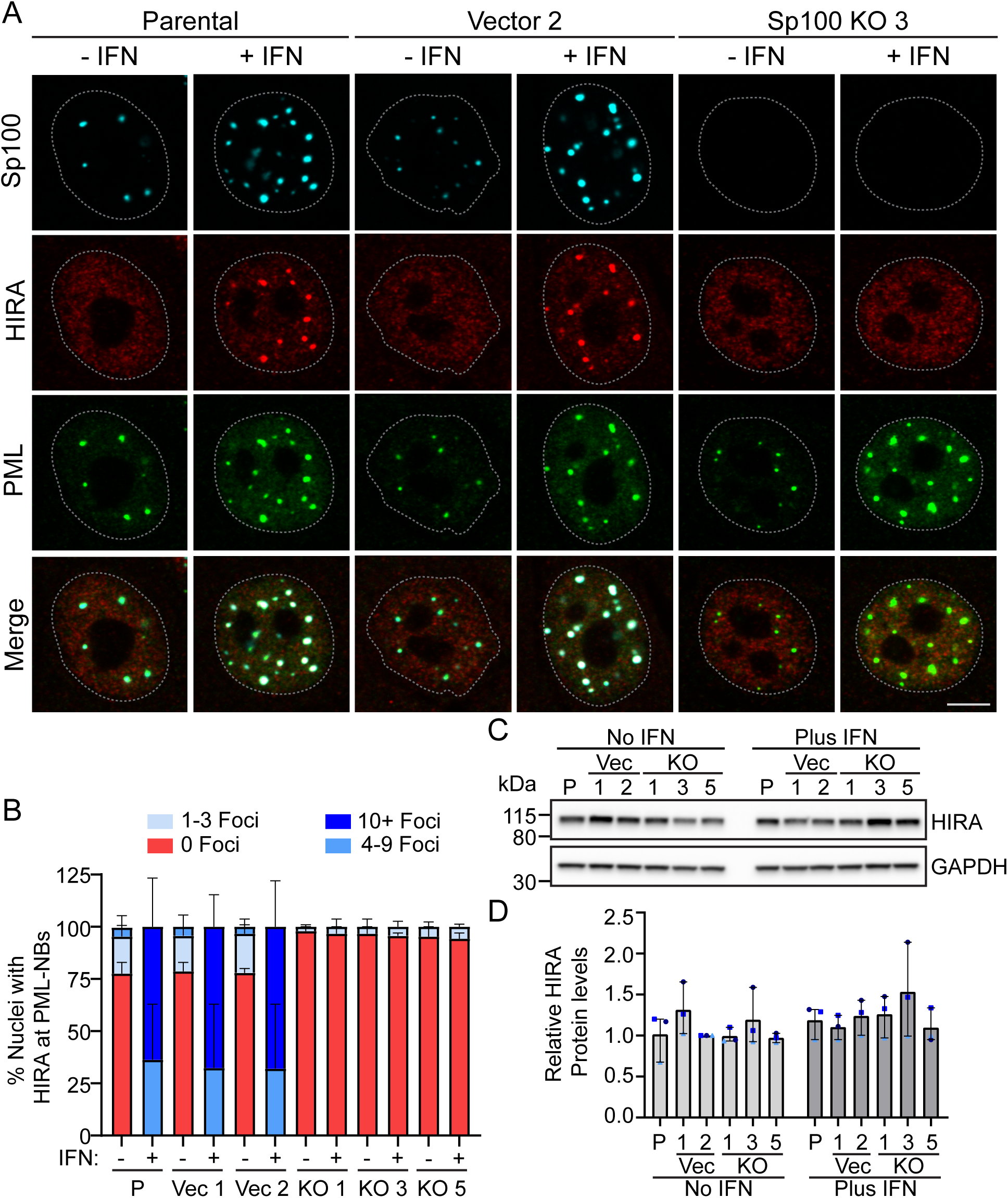
HIRA localization to PML-NBs is dependent on Sp100. **A.** Representative immunofluorescence staining of PML (green), HIRA (red), and Sp100 (cyan) in NIKS Parental, Vector clone 2, and Sp100 KO3 clone cells. Cells were treated with or without IFNα for 48 hours. Data obtained without IFN are shown in Supplementary Figure 4. The white dotted lines outline the nuclei as defined by DAPI staining. Scale bar, 5 μm. **B.** The percentage of nuclei where HIRA was associated with PML-NBs were manually scored in 30 nuclei per experiment (n=3). PML and Sp100 were visualized with PML (sc-9863)/Sp100 (HPA016380) primary antibodies (n=1) or PML (sc-5621)/Sp100 (PCRP-Sp100-1B9-s) primary antibodies (n=2). **C.** Representative immunoblots of HIRA and GAPDH in cell lysates from NIKS Parental, Vector (Clones 1 and 2), and Sp100 KO (Clones 1, 3, and 5) cells treated +/- IFNα for 48 hours. Blots are representative of three independent experiments. **D.** Quantitation of panel C, showing the protein levels of HIRA normalized to GAPDH levels and displayed relative to NIKS Vector 2 minus IFN (n=3). Light, medium, and dark blue shapes denote three independent experiments. All graphs display the mean values from three independent experiments; Error bars represent the range.

To assess whether HIRA protein levels changed by Sp100 knockout, protein lysates prepared from NIKS, vector, and Sp100 KO cells treated with or without type I IFN were analyzed by immunoblot. As shown in Figure 3C and D, HIRA protein levels were unaltered by IFN treatment and were constant among cells with or without Sp100. Taken together, these data show that HIRA protein levels are not changed by IFN treatment or the presence of Sp100 proteins. Rather, Sp100 is required for re-localization of HIRA to PML-NBs and this increased greatly after stimulation with IFN in NIKS keratinocytes.

### Sp100 is required for localization of HIRA complex members to PML-NBs

H3.3 is deposited onto DNA by the HIRA histone chaperone complex that consists of HIRA in association with UBN1 and CABIN1, and which transiently interacts with ASF1a to facilitate histone deposition (Choi *et al*., 2024). Others have shown that all members of the HIRA complex, in addition to ASF1a, localize to PML-NBs in response to various stimuli (Banumathy *et al*., 2009; Kleijwegt *et al*., 2023; Rai *et al*., 2017; Rai *et al*, 2011; Zhang *et al*., 2005). As shown in Figure 3 and by others (Kleijwegt *et al*., 2023; McFarlane *et al*., 2019), HIRA localization to PML-NBs occurs in a Sp100-dependent manner. However, the dependency on Sp100 for localization of other HIRA complex members is unknown. To evaluate this, we assessed the colocalization of HIRA and UBN1 (Figure 4A) or HIRA and ASF1a (Figure 4D) with PML-NBs in Vector 2 and Sp100 KO3 cells, before and after IFN treatment. As shown in Figure 4A, both HIRA and UBN1 localized diffusely throughout the nucleoplasm in Vector 2 cells, with minimal association with PML-NBs in the absence of IFN treatment (Figure 4B). However, they were frequently observed together at PML-NBs following stimulation with IFN. In contrast, neither factor associated with PML-NBs in the Sp100 KO3 cells with or without IFN treatment (Figure 4B). To assess whether protein levels of UBN1 changed with IFN treatment or Sp100 knockout, protein lysates prepared from NIKS Vector 2 and Sp100 KO3 cells treated with or without type I IFN were analyzed by immunoblot. As shown in Figure 4C, UBN1 protein levels were only modestly increased by stimulation with IFN and were constant between cells with and without Sp100.

**Figure 4.**
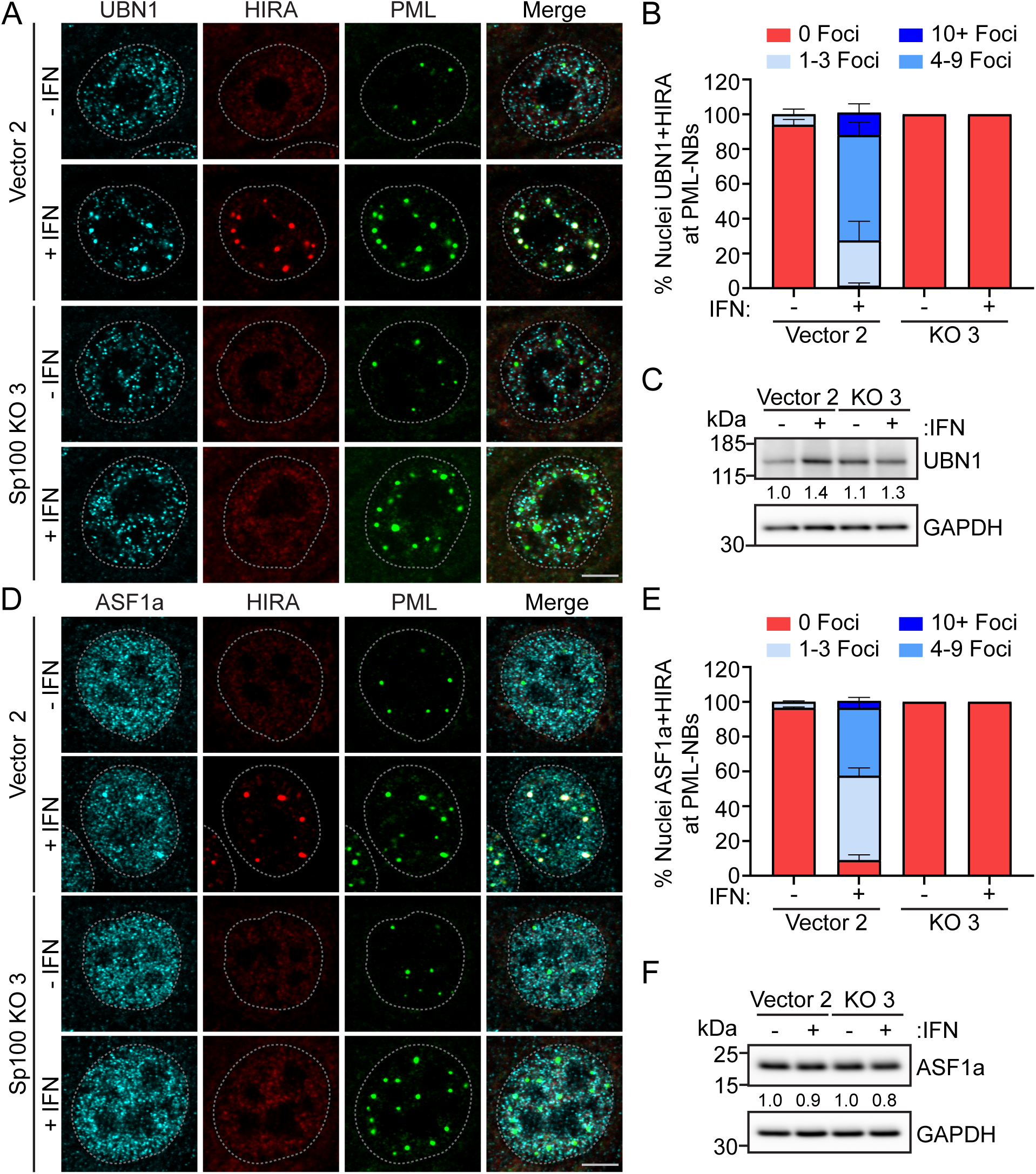
UBN1 and ASF1a localization to PML-NBs occurs in a Sp100-dependent manner. Representative immunofluorescence staining of UBN1 (A) or ASF1a (D) (pseudo-colored cyan), PML (pseudo-colored green), and HIRA (pseudo-colored red) in NIKS vector clone 2 and Sp100 KO3 clone cells with and without IFNα treatment for 48 hours from two independent experiments. A white dotted lines outline the nuclei as defined by DAPI staining. Scale bar, 5 μm. **B and D.** The percentage of nuclei where UBN1 and HIRA (B) or ASF1a and HIRA (E) were associated with PML in +/- IFN conditions. Categories shown are 0 PML-NBs (red), 1-3 PML-NBs (light blue), 4-9 PML-NBs (medium blue), or 10+ PML-NBs (dark blue). For panel B and D quantitation, n = 2; manually scored a minimum of 30 nuclei per experiment. Shown are the mean values of two independent experiments and error bars represent the range. Representative immunoblots of UBN1 (C) or ASF1a (F) and GAPDH in cell lysates from NIKS Vector (Clone 2), and Sp100 KO (Clone 3) cells treated +/- IFNα for 48 hours. Blots are representative of two independent experiments. Average UBN1 or ASF1a levels normalized to GAPDH, relative to Vector 2 – IFN cells are reported below each band.

HIRA and ASF1a were also observed in a nuclear diffuse pattern in NIKS Vector 2 cells without stimulation with IFN (Figure 4D). Stimulation with IFN resulted in frequent association of both HIRA and ASF1a at PML-NBs (Figure 4E). However, similar to HIRA and UBN1, HIRA and ASF1a association with PML-NBs did not occur in Sp100 KO3 cells (Figure 4E). The protein levels of ASF1a were unaltered by stimulation with IFN or by the absence of Sp100 (Figure 4F).

Taken together, these data demonstrate that similar to HIRA (Figure 3C-D), UBN1 and ASF1a protein levels are not greatly changed by IFN treatment or the presence of Sp100 proteins. Instead, localization of HIRA and associated complex members occurs in a Sp100-dependent manner in NIKS keratinocyte cells and increases following stimulation with IFN in Sp100 positive cells.

### The Sp100A isoform promotes HIRA localization to PML-NBs in NIKS Sp100 KO cells

Sp100A, Sp100B, Sp100C, and Sp100HMG are the four main isoforms encoded by alternatively spliced transcripts of the Sp100 gene (Figure 1A). All four isoforms share N-terminal sequences with Sp100A, but Sp100B, Sp100C, and Sp100HMG proteins contain additional domains within the C-terminus that confer the ability of Sp100 to interact with DNA and specific modifications on chromatin (Fraschilla & Jeffrey, 2020). The N-terminal region of all isoforms contains a dimerization region that could promote the formation of homo- and hetero-dimers. We and others have shown that exogenous expression of each isoform results in differential association with PML-NBs and localization patterns within the nucleus (Berscheminski *et al*., 2014; Seeler *et al*, 2001; Stepp, 2015). However, these experiments were performed in cell lines containing endogenous Sp100 proteins that could affect the distribution of exogenous Sp100 proteins. As such, we evaluated the nuclear localization of the four Sp100 isoforms, in addition to a mutated isoform of Sp100B containing a W655Q amino acid substitution in the SAND domain (Sp100BQ) in Sp100 KO3 keratinocytes.

Sp100 KO3 cells were transfected with either a control EYFP expression vector or expression vectors encoding individual Sp100 isoforms as YFP-fusion proteins (Newhart *et al*., 2013). Representative immunofluorescence images of transfected cells treated with or without interferon are shown in Figure 5A, and Supplementary Figure 4, respectively, and the protein levels of each isoform were confirmed by Western blot analysis (Supplementary Figure 5). As shown in Figure 5A and B, all four Sp100 isoforms localized to PML-NBs, but with varied frequencies. Sp100A was most frequently associated with PML-NBs both in the absence or presence of IFN, and Sp100HMG was often associated with PML-NBs. However, Sp100B and Sp100C localized with only with a few PML-NBs per nuclei. It has been shown previously that mutation of the SAND domain in Sp100B results in its re-localization, predominately to PML-NBs (Negorev *et al*, 2006), and our observations confirm this. Stimulation with IFN increased association of each Sp100 isoform with PML-NBs. In addition to localization at PML-NBs, we also observed small patches of Sp100 nuclear accumulation or aggregation of the longer isoforms that did not colocalize with PML, as others have reported (Berscheminski *et al*., 2014; Seeler *et al*., 2001). Thus, individual Sp100 isoforms differentially associate with PML-NBs in the absence of endogenous Sp100 protein.

**Figure 5.**
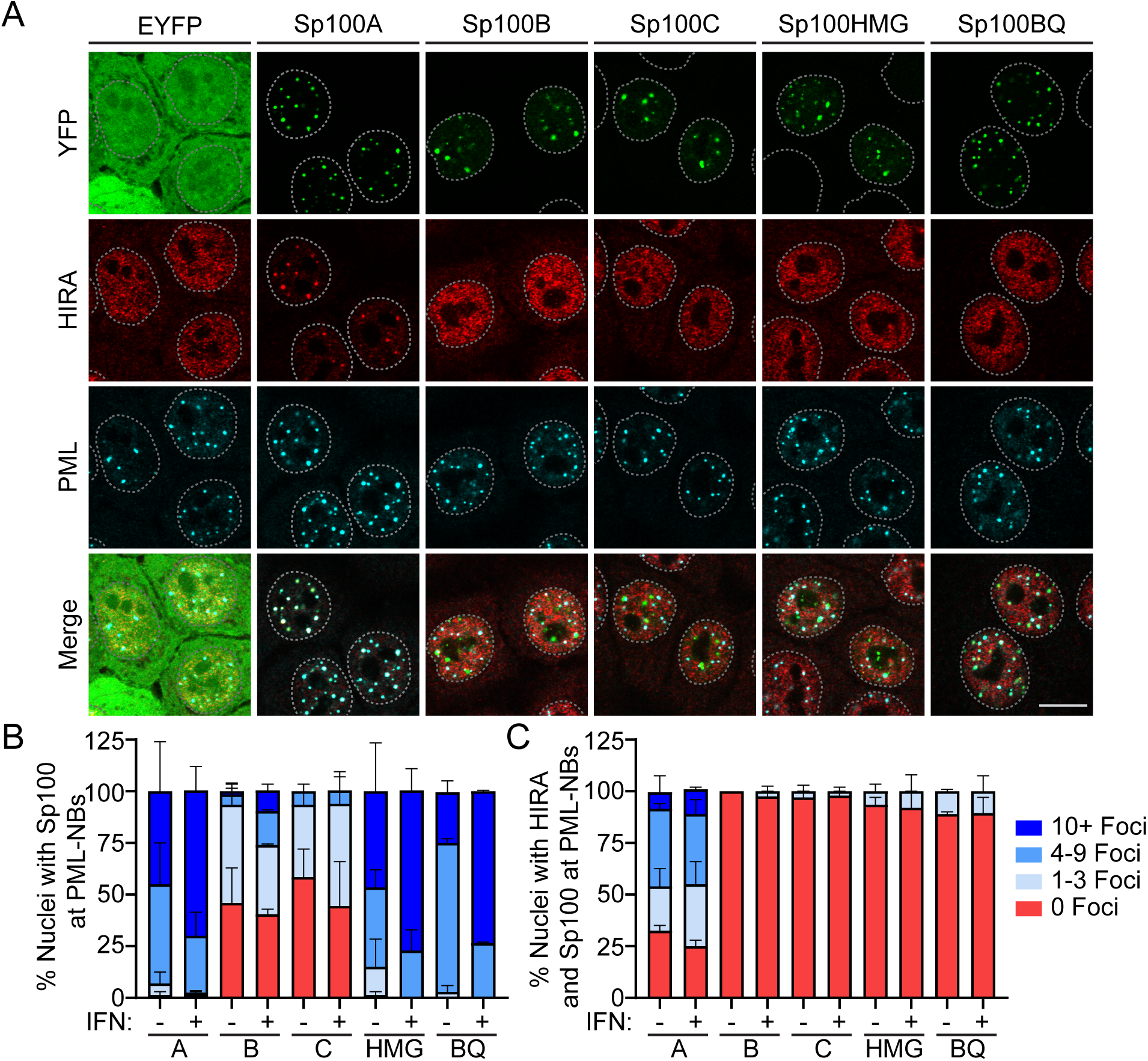
Sp100A promotes HIRA localization at PML-NBs. NIKS Sp100 KO3 cells were transfected with 400 ng of EYFP or EYFP-Sp100 isoforms expression vectors and at 18 hours post-transfection were treated with or without IFNα for 48 hours before fixation. **A.** Representative immunofluorescence staining of YFP (green), HIRA (red), and PML (cyan) in Sp100 KO3 cells (two independent experiments). The white dotted lines outline the nuclei as defined by DAPI staining. Scale bar, 5 μm. Images shown were collected at optimal collection parameters and so are not directly comparable. **B-C.** The percentage of nuclei where Sp100 (B) or HIRA (C) was associated with PML in +/- IFN conditions. Categories shown are 0 PML-NBs (red), 1-3 PML-NBs (light blue), 4-9 PML-NBs (medium blue), or 10+ PML-NBs (dark blue). For panel B and C quantitation, n = 2; manually scored a minimum of 30 nuclei per condition, per experiment. Panel C quantitation is a subset of Panel B; HIRA association with PML-NBs was evaluated in nuclei where the various Sp100 isoforms localized with PML-NBs. Shown are the mean values from two independent experiments and the error bars represent the range.

We next evaluated which Sp100 isoform was responsible for re-localization of HIRA to PML-NBs. Only expression of Sp100A resulted in a high frequency of HIRA and Sp100 colocalization at PML-NBs. This colocalization increased following stimulation with IFN, concomitant with increased association of Sp100A at with PML-NBs (Figure 5B). Despite frequent association of Sp100HMG with PML-NBs (Figure 5B), HIRA was rarely associated with PML-NBs. When Sp100B and Sp100C were associated with PML-NBs, HIRA association with PML-NBs was almost never observed. As shown in Supplementary figure 5, exogenous expression of the Sp100 isoforms did not greatly alter HIRA or PML protein levels. As described above, the Sp100B protein with a mutation in the SAND DNA binding domain localizes primarily to PML-NBs. Despite this, Sp100BQ expression resulted in minimal re-localization of HIRA to PML-NBs. Of note, Sp100A is expressed at higher levels than the other isoforms (Supplementary Figure 5), but examination of Sp100 YFP intensity at individual PML-NBs indicated that HIRA association did not always correlate directly with YFP-Sp100 levels. We conclude that Sp100A is primarily responsible for the re-localization of the histone chaperone HIRA to PML-NBs.

Each Sp100 isoform contains the same N-terminal 483 amino acids, yet only Sp100A protein causes HIRA recruitment to PML-NBs. To investigate this further, we used ColabFold to generate predicted homo-dimeric structures for each of the main Sp100 isoforms (Mirdita *et al*, 2022). As shown in Supplementary Figure 6, the N-terminal region of Sp100A, which corresponds to the previously described HSR domain (Sternsdorf *et al*, 1999), forms a dimeric domain consisting of alpha-helical bundles. The remaining C-terminal residues of Sp100A are predicted to be unstructured. Each of the longer isoforms contained additional structured domains corresponding to the SAND domain, PHD-bromodomain and HMG domain. Each of these domains is predicted to interact with the N-terminal HSR region and could sterically hinder interactions that are important for Sp100A re-localization of HIRA. Each isoform is also predicted to contribute additional disordered sequences that could also mask Sp100A interactions (Supplementary Figure 6).

### Sp100A SUMO interacting motif is important for HIRA localization at PML

To determine what functions of Sp100A are required for its association with PML-NBs and promotion of HIRA localization to PML-NBs, we generated a series of expression vectors that encoded Sp100A proteins mutated in previously characterized functional domains, as shown in Figure 6A (Cuchet-Lourenco *et al*, 2011; Dong *et al*, 2022; Seeler *et al*., 1998; Sternsdorf *et al*., 1999). The mutations targeted the SUMO interacting motif (mSIM) (Cuchet-Lourenco *et al*., 2011), the site of SUMOylation (mSUMO), the nuclear localization signal (mNLS), the HSR domain (ΔHSR) which is required for nuclear body targeting and dimerization (Sternsdorf *et al*., 1999), the HP1 binding region (mHP1) (Seeler *et al*., 1998), alanine and aspartic acid substitutions of the S180 phosphorylation site (Dong *et al*., 2022), and deletion of the last 3 amino acids (ΔKED) that are unique to the Sp100A isoform. Each mutation was substituted into the EYFP-Sp100A expression vector.

**Figure 6.**
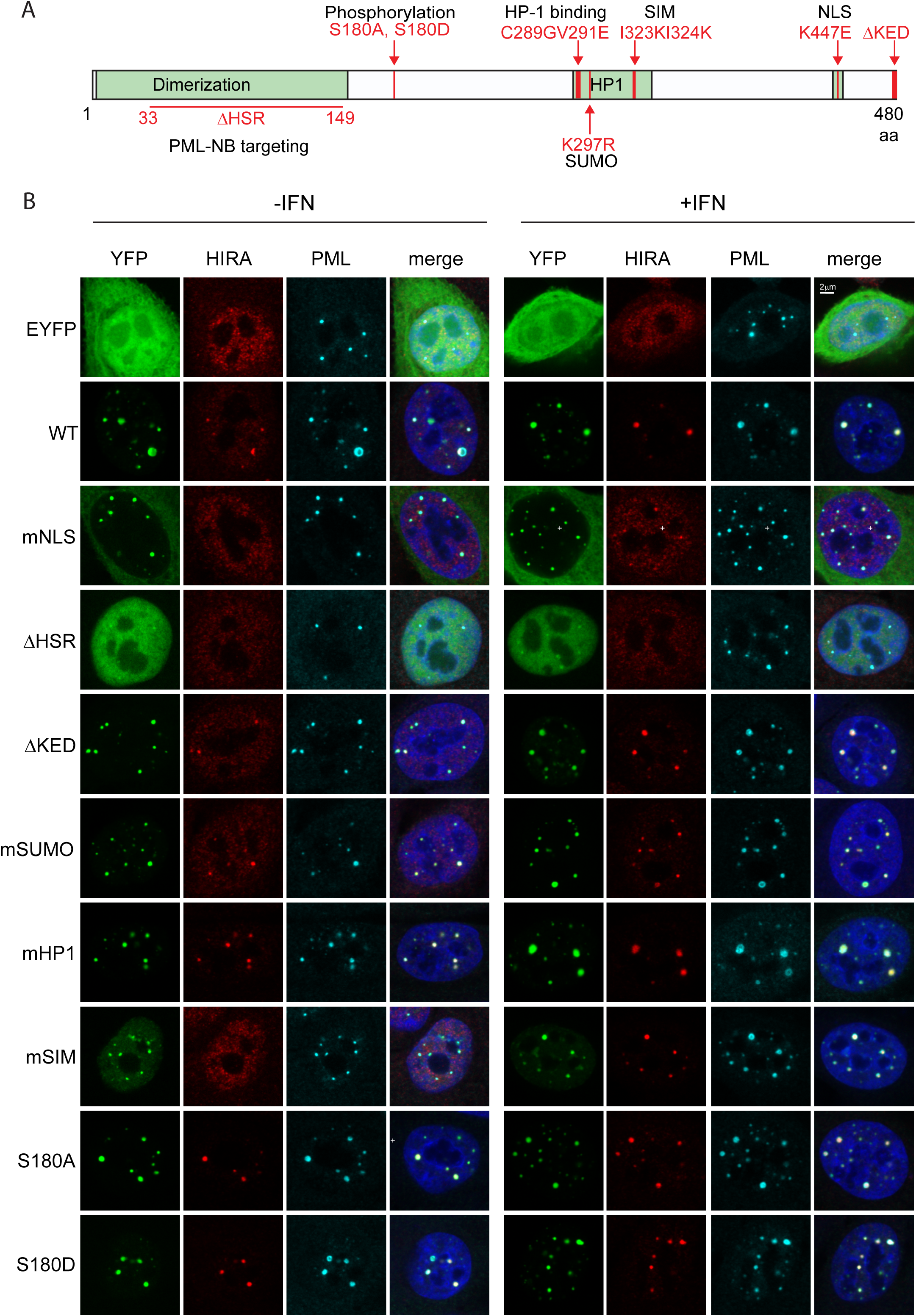
Ability of mutated Sp100A proteins to promote HIRA localization to PML-NBs. **A.** Diagram of Sp100A protein with mutations used in this study indicated. **B.** Representative immunofluorescence staining of YFP-Sp100A (green), HIRA (red), and PML (cyan) in Sp100 KO3 clone cells. Cells were treated with or without IFNα for 24 hours.

To assess the intracellular localization of the mutated EYFP-Sp100A proteins, the expression vectors were transfected into Sp100 KO3 cells. Transfected cells were either untreated or treated with 25ng/ml IFNα at 24 hours post-transfection and fixed 24 hours post-stimulation with IFN. Cells were analyzed by immunofluorescence and confocal microscopy for Sp100 and HIRA intracellular localization (Figure 6B). As shown in the representative images in Figure 6B, and summarized in the Table in Figure 7A, most mutated proteins localized to the PML-NBs similar to the wildtype Sp100A protein (mSUMO, mHP1, and S180 phosphorylation mutations). As expected, the EYFP protein localized throughout the cell, Sp100A ΔHSR localized throughout the nucleoplasm, Sp100A mSIM showed both accumulation at PML bodies as well as a nucleoplasmic distribution, and Sp100A mNLS was mostly cytoplasmic with a small amount entering the nucleus and localizing to PML-NBs. IFN treatment increased the number and amount of PML at PML-NBs, but it did not change the overall intracellular distribution of each Sp100A protein.

**Figure 7.**
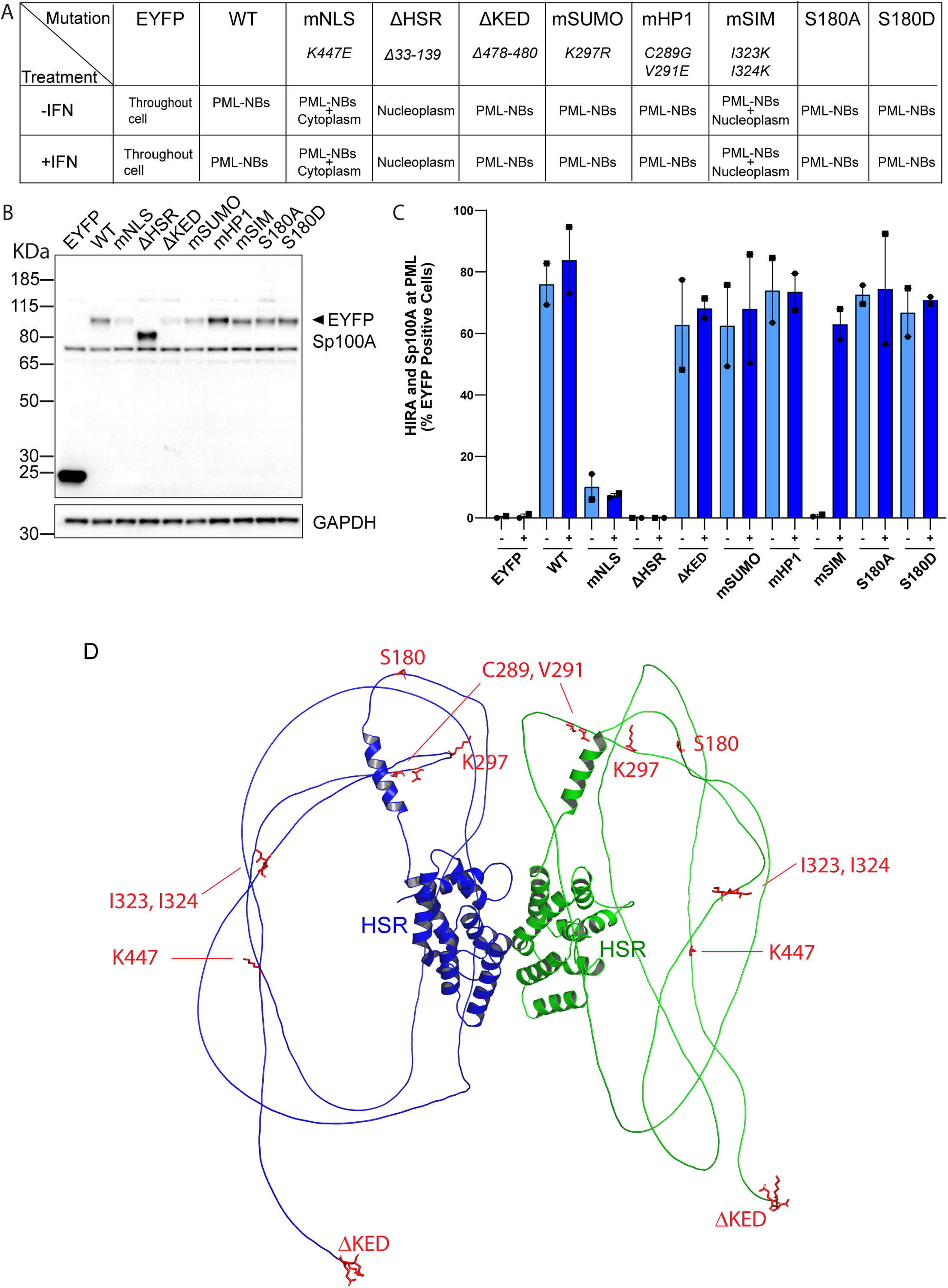
Sp100A protein intracellular location and promotion of HIRA localization to PML-NBs. **A.** Table summarizing the predominant phenotype (in >80% cells) of each YFP-Sp100A mutated protein in Sp100 KO3 cells before and after IFN treatment. **B.** Immunoblot analysis of YFP-Sp100A mutated proteins in NIKS cells. **C.** Quantification of HIRA and Sp100A localization to PML-NBs. Shown are the mean values of two independent experiments with yes/no scoring of at least 70 cells each, and the error bars represent the range. Shapes represent individual experiments. **D**. Structure of homodimer of Sp100A was predicted using Google ColabFold v1.5.5: AlphaFold2 using MMseqs2. The structural HSR domains indicated had high prediction scores and the unstructured regions had low scores. Structures were rendered and mutations highlighted and colored with Pymol (PyMOL Molecular Graphics System, Version 3.0 Schrödinger, LLC).

To confirm that the mutated EYFP-Sp100A proteins were expressed at similar levels and were of the expected molecular weight, the expression vectors were transfected into NIKS cells and Sp100 protein levels analyzed by immunoblot (Figure 7B). Each protein was of the expected size, and all were expressed at comparable levels (mNLS, mSUMO and ΔKED were lower).

To assess the impact of each Sp100A mutation on HIRA localization to PML-NBs, HIRA localization was evaluated in cells with moderate expression of the EYFP-Sp100A proteins (Figure 6B) and is displayed as % cells with HIRA and Sp100A at PML-NBs in Figure 7C. As expected, HIRA did not localize to PML-NBs in NIKS Sp100 KO3 cells expressing YFP alone, even after interferon treatment (Figure 7C). HIRA localized to PML-NBs with Sp100A wildtype, ΔKED, mSUMO, mHP1, S180A and S180D in the majority of cells with or without IFN treatment. Of the remaining mutated Sp100A proteins, for the most part ΔHSR did not localize to PML-NBs and correspondingly did not promote HIRA localization to the bodies. (We do observe a very small amount of the ΔHSR protein enriched in PML-NBs but with no enrichment of HIRA.) Most of the mNLS Sp100A protein was cytoplasmic but small amounts colocalized with PML-NBs, and in 7-10 % these cells HIRA colocalized at PML-NBs. Sternsdorf and colleages have shown that the K447E NLS mutation also greatly reduces SUMOylation of Sp100A (Sternsdorf *et al*., 1999) and this could explain the low efficiency of Sp100-dependent HIRA localization at PML-NBs. However, we and other to find that Sp100A with a mutation in the SUMO site localizes similar to wildtype indicating again that this cannot be the complete explanation for the mNLS phenotype. Most notable is the mSIM Sp100A protein, which localizes to the PML-NBs with or without IFN treatment (there is also a low level of nucleoplasmic signal), but only promoted HIRA colocalization after IFN treatment. This implies that IFN treatment is modifying Sp100A in some way that enables it to promote HIRA colocalization, or alternatively is modifying or inducing other proteins to localize to PML-NBs and co-operate with the mSIM protein to rescue HIRA localization. These results indicate that while the SUMO interacting motif is important, SIM alone is not entirely responsible for the Sp100-dependent HIRA recruitment to PML-NBs

To examine the Sp100A mutations further, they were mapped onto the dimeric Sp100A structure predicted by Colabfold (Figure 7D). With the exception of the ΔHSR mutation that removes most of the structured dimerization and PML-NB targeting regions, all other mutations mapped to the C-terminal regions of the protein predicted to be unstructured and disordered.

### RNA-seq analysis shows no significant changes in basal or interferon induced gene expression in the Sp100 KO cells

To investigate changes in the transcriptome of Sp100 KO cells, and their response to IFN treatment, RNA was collected in triplicate from Vector 2 and Sp100 KO5 cells that were untreated, or treated with IFN for 12 or 48 hours, Figure 8A. Principal component analysis (PCA) of RNA-seq data showed similar basal transcription profiles between Vector 2 and Sp100 KO5 cells, with samples clustering based on interferon treatments rather than genotype (Figure 8B). Minor differences in expression of keratinocyte differentiation markers or interferon-stimulated genes (ISGs) are due to NIKS cell clone-to-clone variation (unpublished data). IFN treatment of both cell lines resulted in a massive induction of interferon inducible genes, but there was no significant difference in the response between the Sp100 positive Vector 2 cells and Sp100 deficient KO5 cells (Figure 8C). We describe above that HIRA will not localize to PML-NBs in the absence of Sp100, and so our findings reinforce those of McFarlane et al. and Kleijwegt et al., who showed that the presence of HIRA at PML-NBs does not change ISG expression (Kleijwegt *et al*., 2023; McFarlane *et al*., 2019).

**Figure 8.**
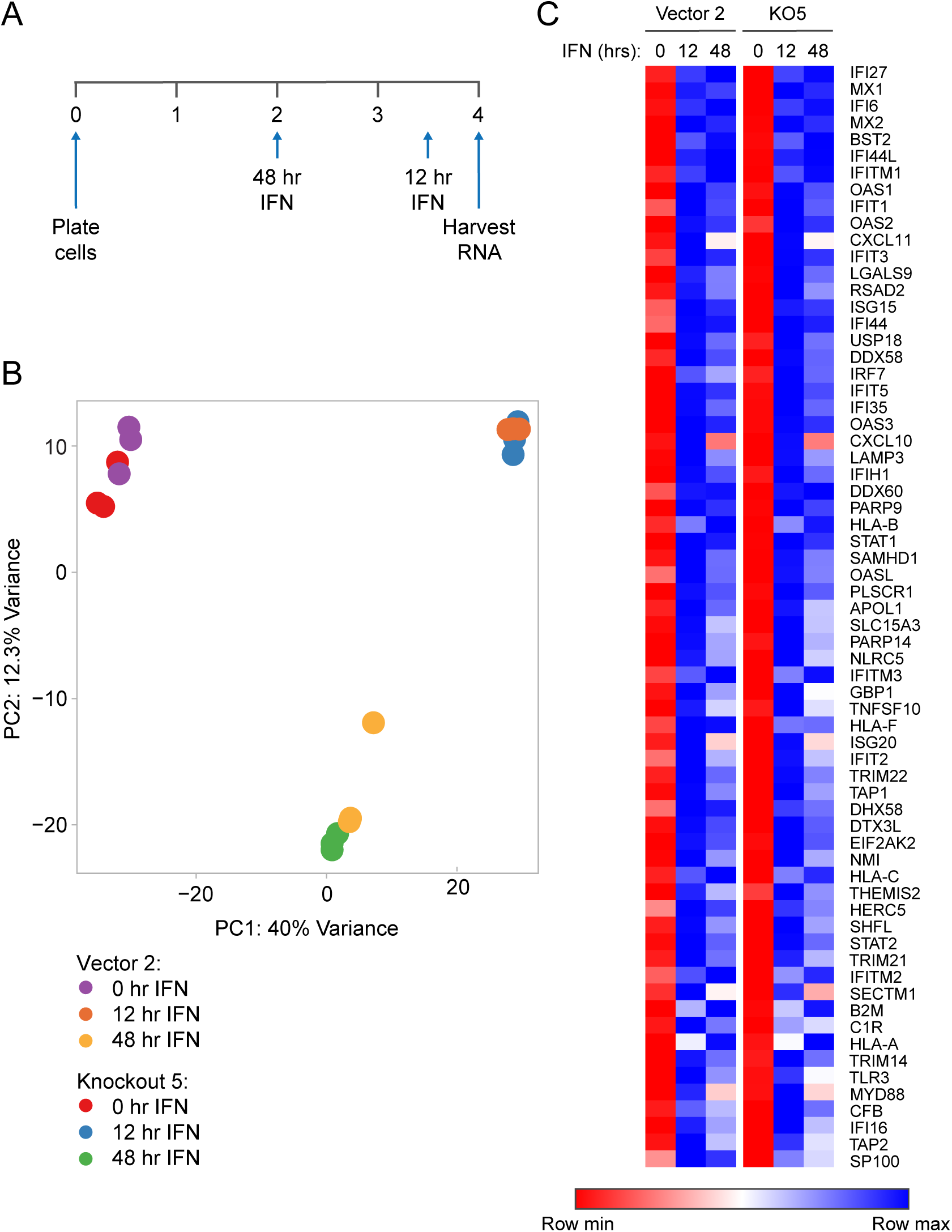
Sp100 KO does not affect IFN stimulated gene expression. RNA-seq analysis was performed in NIKS vector 2 and Sp100 KO5 cells (three replicates per condition). **A.** Experimental timeline: RNA was isolated from untreated cells or cells treated for 12 or 48 hours with IFNα. **B.** Principal component analysis to show variance in gene expression across the RNA-seq samples. Colored circles represent individual replicates for the indicated conditions. PC, Principal component. **C.** Heatmap of genes identified by K-means clustering analysis to be enriched in the immune response pathway. Data is represented as the mean value across the three replicates for each condition. Each row in the heatmap is color-scaled independently, with colors representing the row’s minimum to maximum value. Of note, low level RNA-seq reads were detected for the Sp100 transcripts in the Sp100 KO5 cells, but no protein is detected (Supplementary Figure 1).

## Discussion

In this study we generated CRISPR Sp100 KO cell lines from the immortalized keratinocyte cell line, NIKS, and showed that they could differentiate into a stratified epithelia similar to the parental NIKS cells. The absence of Sp100 did not perturb PML-NB formation nor recruitment of Daxx to PML-NBs. Recent studies have shown that the recruitment of the histone chaperone HIRA to PML-NBs is dependent on the Sp100 protein (Kleijwegt *et al*., 2023; McFarlane *et al*., 2019). Thus, the NIKS Sp100 KO cell lines are valuable to study the interplay of HIRA and Sp100 in human keratinocytes.

The histone chaperone, HIRA localizes to PML-NBs in response to viral infection or transfection with plasmid DNA (Cohen *et al*., 2018; McFarlane *et al*., 2019; Rai *et al*., 2017), both of which induce secretion of IFNs (Li *et al*, 2005; MacMicking, 2012). Similar to previous studies, we observed that IFN treatment caused re-localization of HIRA to PML-NBs in NIKS keratinocytes and this is independent of viral infection or cellular senescence (Kleijwegt *et al*., 2023; McFarlane *et al*., 2019). We show that HIRA (and other components of the chaperone complex, Ubn1 and Asf1) do not localize to PML-NBs in NIKS keratinocytes in the absence of Sp100, even in the presence of interferon.

The Sp100 KO NIKs cells present a unique opportunity to evaluate the roles of the different Sp100 isoforms in the absence of endogenous Sp100 proteins. Previous studies have shown that each isoform localizes with varying degrees to PML-NBs in cells that express endogenous Sp100 (Berscheminski *et al*., 2014; Seeler *et al*., 2001; Stepp, 2015). In the Sp100 KO keratinocytes, Sp100A isoform associates predominately with PML-NBs and the longer isoforms associate partially with PML-NBs. The longer forms contain chromatin and DNA binding domains that can direct their association to other regions of the nucleus and in support of this we show that mutation of the SAND DNA binding function in the Sp100B isoform results in a protein that predominantly localizes with the PML-NBs. We also show that only the Sp100A isoform could re-localize HIRA to PML-NBs. This is surprising because each isoform contains the same amino acid sequences as Sp100A (except for the last three amino acid residues). The Sp100A protein is expressed at higher levels than the other isoforms, but this does not explain why it is the only isoform to re-localize HIRA as the mutated Sp100B W655Q protein localizes strongly to PML-NBs but does not change HIRA localization. It is possible that Sp100A is differentially post-translationally modified compared to the other isoforms, or that interaction between N-terminal and C-terminal Sp100 domains sterically hinder the ability of the longer forms to re-localize HIRA.

We show that Sp100A is the predominant isoform responsible for HIRA localization to PML-NBs. Expression of Sp100A proteins with previously characterized mutations indicates that (as also shown by others (Kleijwegt *et al*., 2023)) mutation of the major SUMO site of Sp100 at K297 does not affect Sp100A localization to PML-NBs nor recruitment of HIRA. Not unexpectedly, deletion of the HSR PML-NB targeting domain abrogates both functions. However, mutation of the SIM motif does not interfere with Sp100A’s PML-NBs localization but does abrogate HIRA recruitment to the nuclear bodies. While this could be postulated to be due to Sp100A recruiting HIRA through SUMO-SIM interactions, Kleijwegt and colleagues have shown that mutation of putative SUMO sites in HIRA residue K809 does not affect its accumulation at PML-NBs (Kleijwegt *et al*., 2023) and no direct protein interaction could be detected between HIRA and Sp100 (McFarlane *et al*., 2019). In contrast, Kleijwegt et al. show that mutation of the SIM2 motif in HIRA (225VLRL228) abrogated its localization to PML-NBs (Kleijwegt *et al*., 2023). In our study, treatment of cells expressing Sp100A mSIM with IFN resulted in localization of HIRA to PML-NBs. IFN treatment results in SUMOylation of many proteins (Sahin *et al*, 2014), particularly at PML-NBs, and we postulate that an increase in SUMOylation greatly enhances the SUMO-SIM interactions among many proteins. It may be possible to dissect these interactions more completely by generating combinatorial mutations in the Sp100A protein.

The concentration of macromolecules at nuclear bodies can greatly increase the efficiency of biochemical processes. PML-NBs are thought to form biomolecular condensates that might be dependent on liquid-liquid phase separation (LLPS) (Alberti *et al*, 2019; Corpet *et al*., 2020). These condensates are driven by multivalent interactions such as those between proteins that are SUMOylated and those with SUMO interaction motifs (SIMs) (Liebl & Hofmann, 2022). Many of these proteins also contain intrinsically-disordered regions (IDRs), which can present flexible interaction motifs to partner proteins (Uversky, 2017). Our structural predictions of the Sp100 isoforms in Figures 7 and Supplementary Figure 6 show that sequences outwith the HSR, SAND, PHD, bromodomain and HMG domains are unstructured and are likely IDRs. Of note, most functional motifs mapped to the Sp100A protein localize to these regions.

Interferon stimulation results in rapid transcriptional activation of thousands of ISGs. These gene loci localize adjacent to PML-NBs after interferon treatment, and HIRA has been shown to deposit H3.3 at the 3’ end of these genes (Kleijwegt *et al*., 2023). However, this deposition occurs well after the rapid induction of ISG transcription, and HIRA localization to PML-NBs is not absolutely required for the IFN response (this study and (Kleijwegt *et al*., 2023; McFarlane *et al*., 2019)). Alternatively, the HIRA-dependent deposition of H3.3 could generate epigenetic memory at ISGs to enable a more rapid response upon repeated stimulation (Kamada *et al*, 2018).

Despite the absence of Sp100 proteins and no localization of HIRA to PML-NBs, the NIKS Sp100 KO keratinocytes described in this study show normal growth and differentiation in cell culture. However, Sp100 proteins have potent pro-viral and anti-viral responses and thus the NIKS Sp100 KO cells will be valuable for the study of viral infection of keratinocytes in monolayer culture or in a stratified epithelium. Furthermore, while the Sp100 KO cells have an efficient immediate response to IFNα treatment, they could prove useful for studying the role of Sp100 and HIRA in long-term epigenetic innate immune memory.

## Materials and Methods

### Cell culture

NIKS cells (RRID:CVCL_A1LW) were grown in Rheinwald-Green F medium (3:1 Ham’s F-12/high-glucose Dulbecco’s modified Eagle’s medium [DMEM], 5% fetal bovine serum [FBS], 0.4 µg/ml hydrocortisone, 8.4 ng/ml cholera toxin, 10 ng/ml epidermal growth factor, 24 µg/ml adenine, 2mM glutamine, 6 µg/ml insulin, 100 U/ml penicillin, and 100 µg/ml streptomycin) on a layer of lethally irradiated J2/3T3 mouse fibroblast cells. J2/3T3 fibroblasts were grown in DMEM, 10% newborn calf serum, 2 mM glutamine,100 U/ml penicillin, and 100 µg/ml streptomycin. Parental NIKS cells were profiled by KaryoStat Karyotyping (Thermo Fischer). All cell lines tested negative for mycoplasma contamination in March 2024.

#### Organotypic rafts

Organotypic raft cultures were generated by plating NIKS keratinocytes on a collagen dermal equivalent containing NIH 3T3 fibroblasts as described previously (Porter *et al*, 2021). Rafts were lifted to the air-liquid interface and cultured for 15 days in raft culture medium (3:1 DMEM/F12, 10% fetal calf serum [FCS], 0.4 μg/ml hydrocortisone, 0.01 nM cholera toxin, 5 μg/ml transferrin). Rafts were fixed in 3.7% formaldehyde, embedded in paraffin and sections stained with hematoxylin and eosin.

#### Generation of Sp100 knockout cell line using CRISPR

Guide RNAs (gRNAs) were designed using CHOPCHOP, https://chopchop.cbu.uib.no/, a webtool for designing target sites for CRISPR Cas9 mutagenesis, and the UCSC Genome Browser (Perez *et al*, 2025) to check for specificity. The top five gRNAs that targeted all isoforms of Sp100 were selected. The oligos used were 5’-AGGATGTTCACGGAAGACCA-3’ (exon 3); 5’-CTTTCTTCATCACAGGGCAACGG-3’ (exon 8), 5’-TGCTGTCCAAGTGAATAATGGGG-3’ (exon 8), 5’-CAAGTGAATAATGGGGATGCTGG-3’ (exon 8) and 5’-ACTCTTTTCGAAGCCTGACTTGG-3’ (exon 6). The oligonucleotides were cloned into pSpCas9(BB)-2A-GFP (PX458, Feng Zhang Addgene plasmid # 48138). The resulting five plasmids were mixed in equal ratios and electroporated into NIKS cells using the Amaxa Human Keratinocyte Nucleofector Kit (Lonza). Vector control cells were electroporated with pSpCas9(BB)-2A-GFP without the gRNA sequence. Positively transfected cells were isolated by fluorescence activated cell sorting (FACS) for GFP expression at 72 hours post-transfection. The GFP positive cells were plated on a 10cm plate of lethally irradiated J2/3T3 fibroblasts and allowed to recover and expand for eight days. Cells were plated at 100 cells per well of a 12-well plate and the resulting cells were screened for the absence of Sp100 by immunofluorescence staining with the Sp100 antibody, rabbit anti-Sp100 HPA016707 (Sigma). Pools that showed no expression of Sp100 by immunofluorescence were expanded into cell lines.

#### Plasmids

YFP-Sp100 isoform expression vectors in pEYFP (pEYFP-Sp100A, pEYFP-Sp100B, pEYFP-Sp100C, pEYFP-Sp100HMG, and pEYFP-Sp100B W655Q) were acquired from Susan Janicki and have been previously described (Newhart *et al*., 2013). To generate mutations in Sp100A, gene fragments containing point mutations or deletions in Sp100A sequences were synthesized by Synbio. The mutated fragments were cloned between the BamHI/Mfel or EcoRI/BamHI sites in pEYFP-Sp100A. The S188D and S188A mutations described by Dong et al. (Dong *et al*., 2022) correspond to serine residue 180 in the sequence of Sp100A used in this study (NP_001193631.1). The sequences were verified by long-read sequencing technology (Plasmidsaurus).

#### Transfection of EYFP-Sp100 Expression Vectors

NIKS Sp100 KO cells were seeded at a density of 2.9 x 10^4^ cells/cm^2^ onto a monolayer of irradiated J2/3T3 cells in a 12-well plate containing glass coverslips or in a 6-well plate and cultured overnight. Cells were transfected with 400 ng control EYFP expression vector or EYFP expression vectors encoding Sp100A, Sp100B, Sp100C, Sp100HMG, Sp100B-W655Q, or Sp100A mutations complexed with 3:1 Fugene6 transfection reagent according to the manufacturer’s instructions. Cells were stimulated with 25 ng/mL Human Interferon Alpha 2a (PBL Assay Science), as indicated. Cells were fixed on coverslips or lysed for protein extraction, as indicated.

#### Immunofluorescence

Cells were fixed at room temperature in 4% paraformaldehyde/1x PBS for 15 minutes, permeabilized with 0.1% or 0.5% Triton-X 100 in PBS, blocked in 5% normal donkey serum, and stained with the following primary antibodies for either 1 hour at 37 ℃ or overnight at 4℃: Sp100 rabbit polyclonal [RRID: AB_1857398] (Sigma; HPA016707, 1:500), Sp100 mouse monoclonal [AB_2722351] (Development Studies Hybridoma Bank; PCRP-SP100-1B9-s, 5 µg/mL), PML [PG-M3] mouse monoclonal [RRID: AB_628162] (Santa Cruz; sc-966, 1:50-1:100), PML [A-20] goat polyclonal [RRID: AB_2284064] (Santa Cruz; sc-9862, 1:50), PML [H-238] rabbit polyclonal [RRID: AB_2166848] (Santa Cruz; sc-5621, 1:100), PML [G-8] mouse monoclonal (Santa Cruz; sc-377340, 1:50), Daxx rabbit polyclonal [RRID: AB_1078625] (Sigma; HPA008736, 1:100), HIRA [WC119] mouse monoclonal [RRID: AB_1977097] (Millipore; 04-1488, 1:100), UBN1 rabbit polyclonal [RRID: AB_2684420] (Sigma; HPA061029), ASF1a [C6E10] rabbit monoclonal [RRID: 2289918] (Cell Signaling; 2990S, 1:100), and GFP-Booster Alexa Fluor 488 alpaca [RRID: 2827573] (Chromotek; gb2AF488-50:50 µl, 1:1,000). Fluorescent secondary antibodies conjugated with Alexa 488, Alexa 594, Rhodamine Red-X, or Alexa 647 (Jackson Immunoresearch; 1:100) were applied using standard procedures. The coverslips were stained with 300nM DAPI and mounted with Prolong Gold.

#### Confocal microscopy

Images were collected with a TCS-SP8 laser scanning confocal imaging system using a 63X oil immersion objective (numerical aperture 1.4). 2D images are from a single optical slice unless otherwise mentioned. All images were processed using Leica LAS AF Lite (Leica Microsystems, version 2.6.0) or Leica Application Suite X (Leica Microsystems, version 3.7.0.20979) software or Imaris (version 10.2, Bitplane). Images were pseudo-colored where indicated.

#### Image analysis

Manual Scoring: The average number of visible PML-NBs per nuclei was counted for a minimum of 30 nuclei per replicate. The percentage of visible PML-NBs per nuclei containing Daxx was assessed for a minimum of 30 nuclei per replicate. The association of various proteins (HIRA, HIRA+UBN1, HIRA+ASF1a, and Sp100) with visible PML-NBs was scored, grouped into four categories according to the number of positive PML-NBs (0 foci, 1-3 foci, 4-9 foci, and 10+ foci), and reported as the total percentage of nuclei associated with each category for a minimum of 30 nuclei per replicate. HIRA association with PML-NBs in cells containing EYFP-Sp100 isoforms was evaluated as described above but, only nuclei where Sp100 was associated with PML-NBs were scored. The sample size per condition is at least two biological replicates. Counts were non-blinded.

For assessment of the mutated Sp100A proteins, cells with moderate YFP expression were identified and the intracellular location of the mutated Sp100A proteins scored. HIRA localization to PML-NBs was assessed by manual counting of HIRA accumulation (yes/no) at PML-NBs in cells with Sp100A localizing to PML-NBs. Between 35 and 250 cells were scored for each condition per replicate. The sample size per condition is at least two biological replicates. Counts were non-blinded.

#### Western blot

Feeders were removed prior to protein extraction. Cells were lysed in SDS extraction buffer (50 mM Tris-HCl [pH 6.8], 5% [wt/vol] SDS, 10% [vol/vol] glycerol), sonicated, and heat denatured. A BCA protein assay kit (Thermo-Pierce) was used to determine total protein concentration. Prior to loading the gel, NuPAGE LDS Sample Buffer© and DTT (50 mM) were added to the protein samples, which were heated at 70°C for 10 minutes. 12-15 µg total protein was separated on 4 to 12% or 8% bis-Tris polyacrylamide gels (Invitrogen) from one or two technical replicates per biological replicate. Proteins were transferred overnight onto 0.45 µm polyvinylidene difluoride membrane (Millipore) and proteins detected by immunoblotting. The antibodies used were Sp100 rabbit polyclonal [RRID: AB_1857398] (Sigma, HPA016707; 1:1,000 or 1:2,000), PML rabbit polyclonal [RRID: AB_873108] (Bethyl; A301-167A, 1:1,000), Daxx rabbit polyclonal [RRID: AB_1078625] (Sigma; HPA008736, 1:1,000), HIRA [WC119] mouse monoclonal [RRID: AB_1977097] (Millipore; 04-1488, 1:500 overnight at 4 ℃), UBN1 rabbit polyclonal [RRID: AB_2684420] (Abcam; ab101282-1001, 1:1,000), ASF1a [C6E10] rabbit monoclonal [RRID: 2289918] (Cell Signaling; 2990S, 1:1,000 overnight at 4 ℃), GFP rabbit polyclonal [RRID: AB_303395] (Abcam; ab290, 1:2,500), GAPDH [6CS] mouse monoclonal [RRID: AB_627679] (Santa Cruz; sc-32233, 1:1,000), and Tubulin [DM1A] mouse monoclonal [RRID: 477583] (Sigma; T6199, 1:1,000). Species-appropriate secondary antibodies conjugated to horseradish peroxidase (Pierce) were used at a 1:10,000 dilution and detected with SuperDura Extended Western Detection Reagent (Thermo-Pierce). The chemiluminescent signal was collected with a Syngene G:Box. Syngene Gene Tools software (version 4.3.9.0 or 4.3.17.0) was used to quantitatively assess protein levels. Raw protein levels were obtained by subtraction of a background region and normalization according to their associated GAPDH level.

#### RNA extraction

Total RNA was isolated with the RNeasy Mini-RNA extraction kit (Qiagen). One microgram of total cellular RNA was reverse transcribed with the Transcriptor First-Strand Synthesis kit (Roche).

#### RNA sequencing (RNA-seq)

Keratinocytes were seeded onto 10cm plates of irradiated J2/3T3 fibroblasts at 3.5x10e4 cells per plate and harvested for RNA four days later. Cells were untreated or stimulated with 25 ng/mL Human Interferon Alpha 2a (PBL Assay Science) for 12 or 48 hours prior to harvesting. Feeders were removed prior to RNA extraction. Total RNA from three replicates was isolated using the RNeasy Mini-RNA extraction kit (Qiagen). RNA concentration and integrity was determined using the Quibit Fluorometer (Invitrogen) and Bioanalyzer 2100 (Agilent Technologies), respectively. Polyadenylated RNA was sequenced (2 x 76 bp paired-end reads) using the Illumina NextSeq (CCR Sequencing Facility, FNLCR/NCI/NIH). RNA-seq data was aligned to the human hg19 reference genome using STAR (version 2.6.1). K-means clustering was performed using iDEP 2.01 (Ge et al, 2018) and heatmaps were generated using Morpheus, https://software.broadinstitute.org/morpheus. RNA-seq datasets are accessible through GEO Series accession number GSE290877.

## Acknowledgments

We thank all members of the McBride laboratory for helpful advice and discussions, Dr. Justin Lack for advice on sequencing and Dr. Tovah Markowitz (RTB/NIAID/NIH) for processing RNA-seq data. RNA sequencing was performed by the CCR (Center for Cancer Research) Sequencing Facility (FNLCR/NCI/NIH). A portion of this manuscript is derived from the thesis of ADF (Della Fera, 2024).

