## Supplemental Table and Figures for "Sp100A isoform promotes HIRA histone chaperone localization to PML nuclear bodies"

**Supplementary Table 1: Genotype of Sp100 CRISPR knockout cells**

| | **Exon 3** | **Exon 6** | **Exon 8** |
|---|---|---|---|
| **CRISPR guide** | 1 | 2 | 3, 4, 5 |
| **Vector 1 #9** | wildtype | wildtype | wildtype |
| **Vector 2 #10** | wildtype | wildtype | wildtype |
| **Sp100 KO 1 #11** | Wildtype, 15bp del1, 15bp del2 | wildtype | 35bp del1, 35bp del2 |
| **Sp100 KO 2 #12** | Wildtype, 11bp del 16bp del | wildtype | 35bp del1, 35bp del2 |
| **Sp100 KO 3 #13** | Wildtype, 15bp del1, 15bp del2 | wildtype | 35bp del1, 35bp del2 |
| **Sp100 KO 4 #28** | 158bp 28S rRNA insertion | wildtype | 35bp del, 36bp del |
| **Sp100 KO 5 #33** | 158bp 28S rRNA insertion | wildtype | 36bp del |

Supplemental figure 1

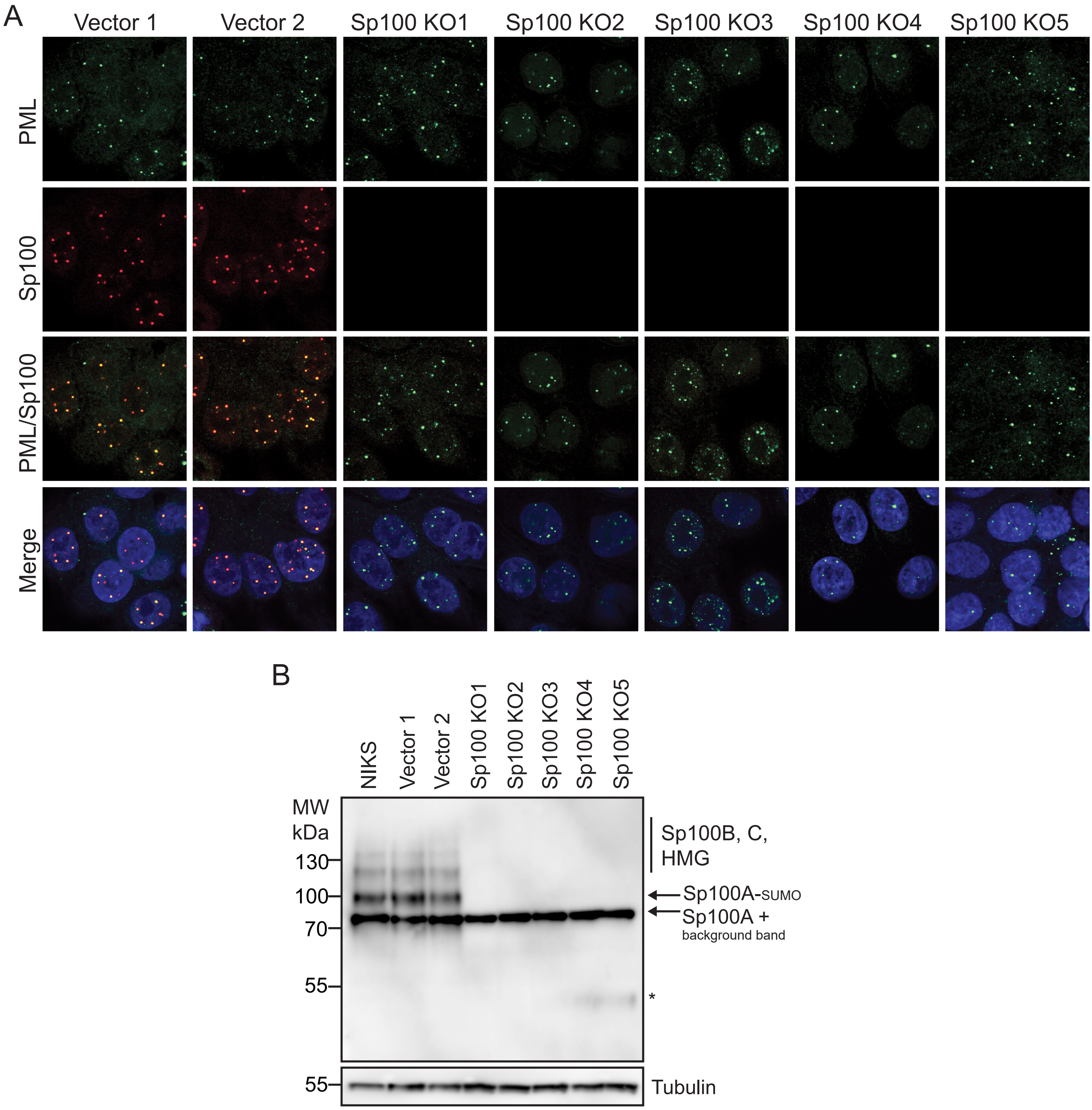

Supplemental figure 2

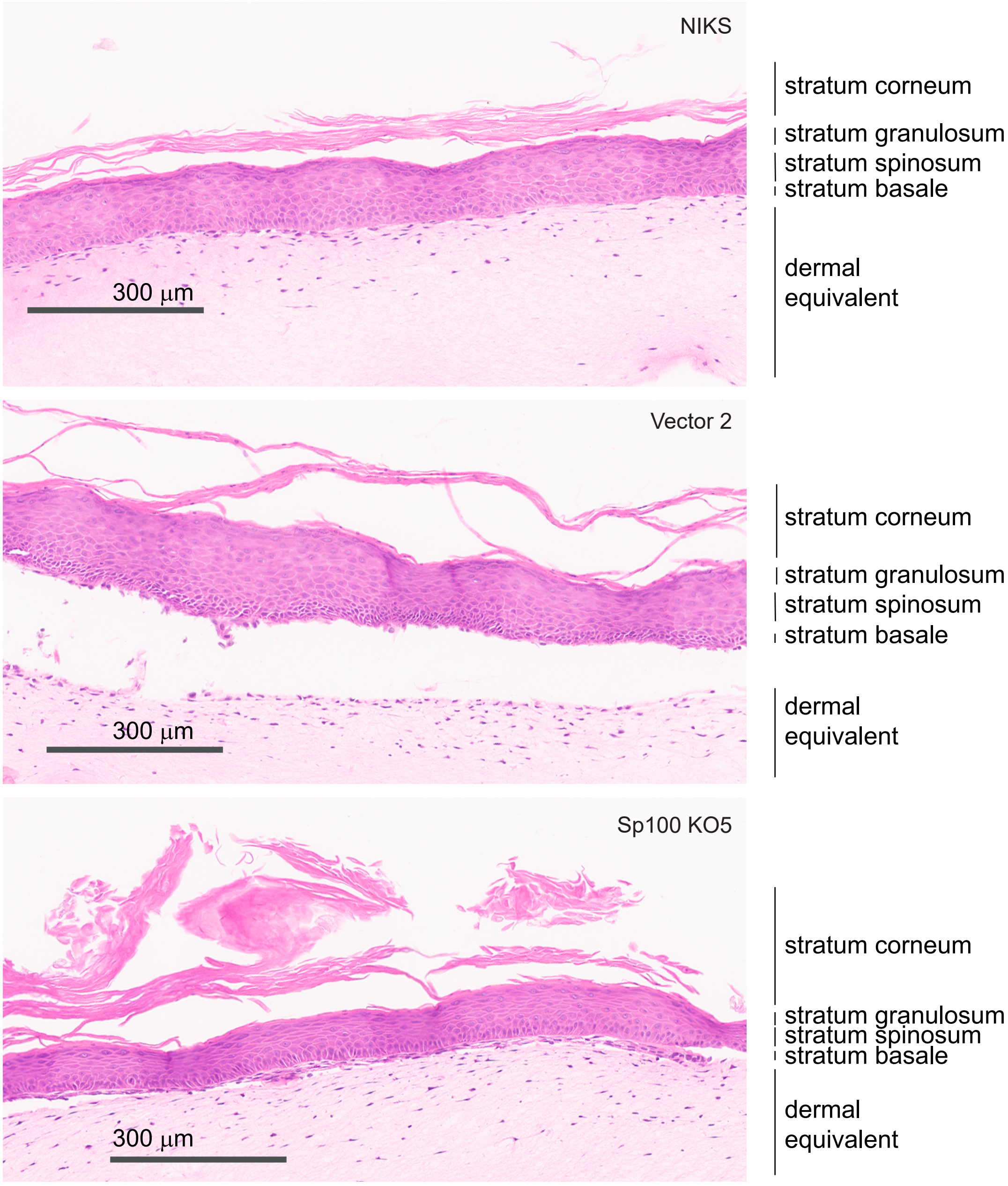

Supplemental figure 3

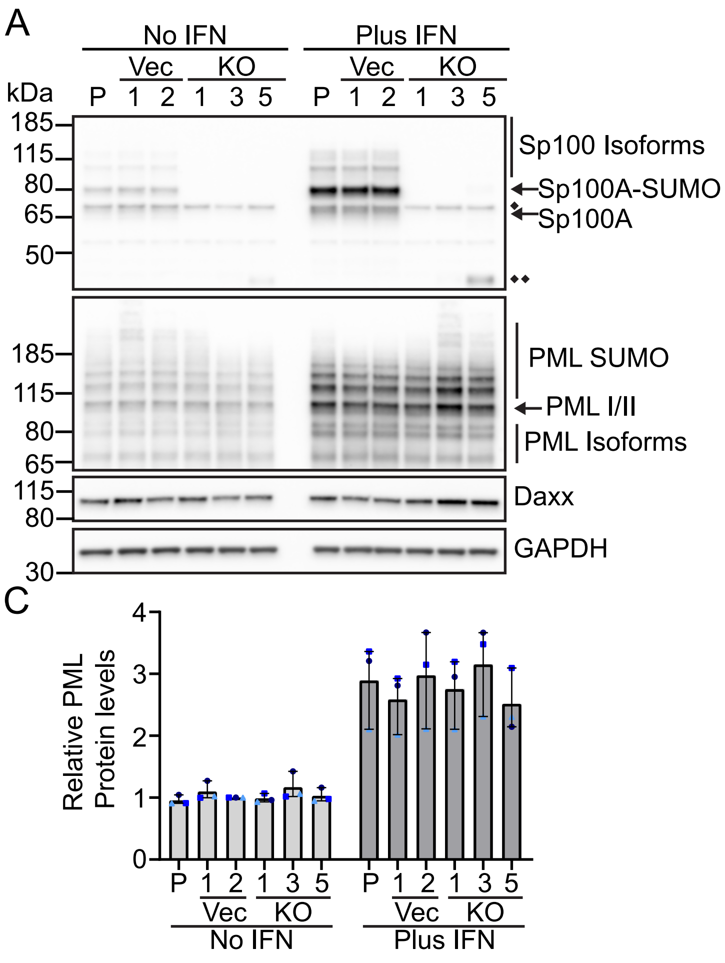

Supplemental figure 4

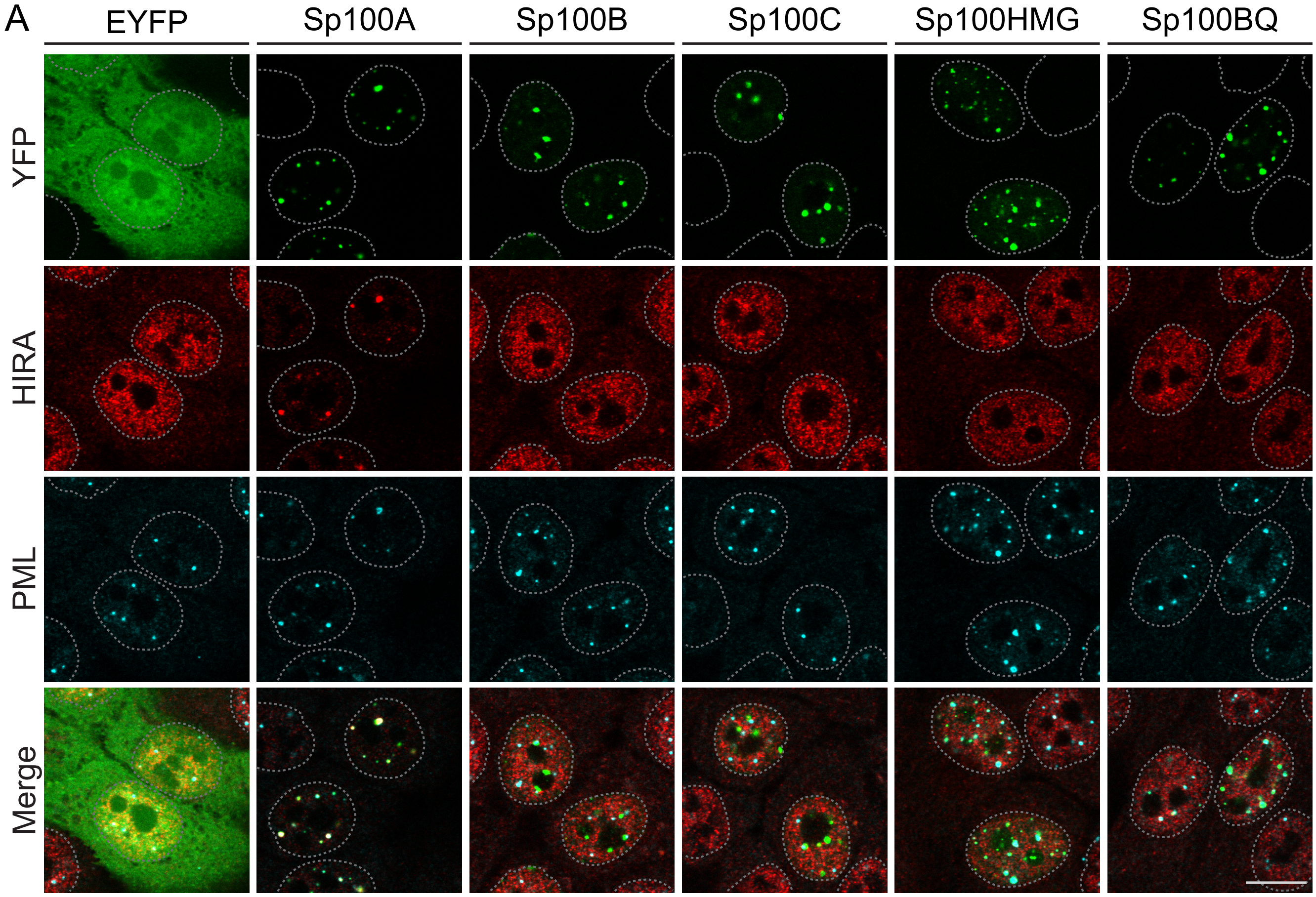

Supplemental figure 5

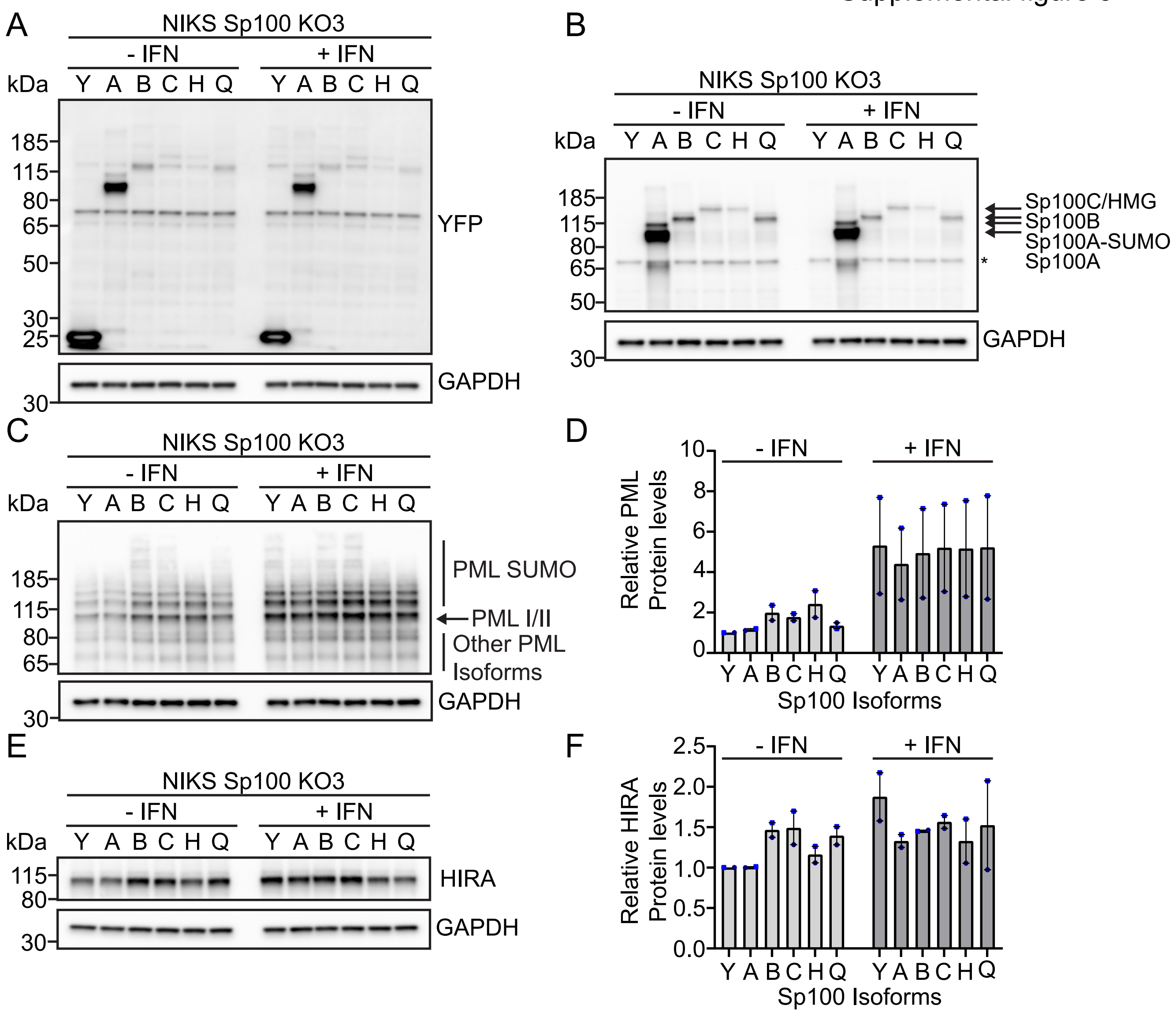

Supplemental figure 6

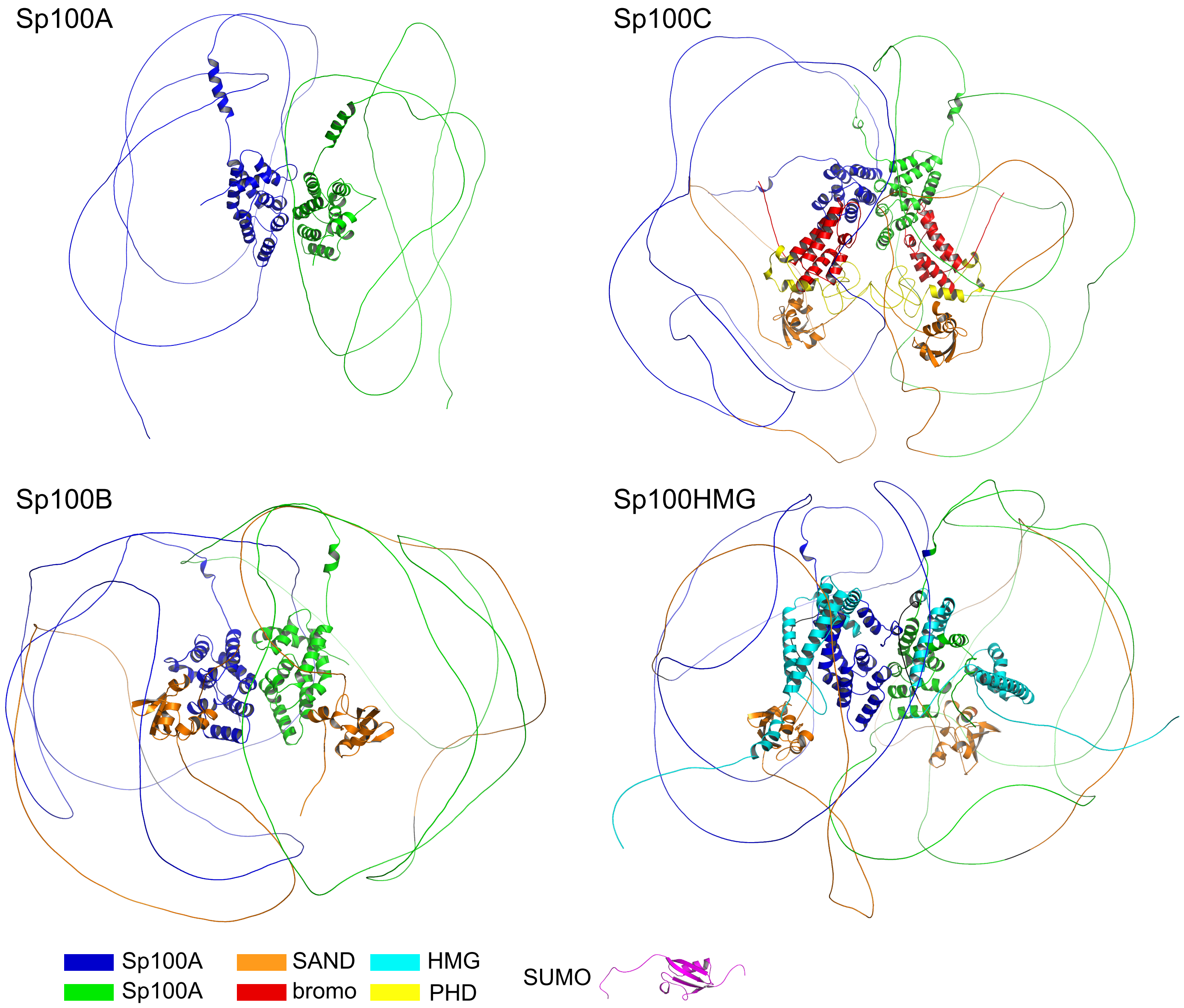
